# Identification of critical cell-types using genetic modules: A case study of neurodevelopmental disorders

**DOI:** 10.1101/2023.07.04.547726

**Authors:** Julie Chow, Marketa Tomkova, Ashleigh Thomas, Elior Rahmani, Sagiv Shifman, Fereydoun Hormozdiari

## Abstract

Identifying the critical cell-types impacted by various diseases is crucial for understanding disease mechanisms and developing targeted therapeutics. Recent advances in disease genetic module discovery and single-cell technologies provide a unique opportunity to study critical cell-types based on functional pathways and modules. Disease genetic modules are defined as sets of genes with correlated expression that are part of the same biological pathways and are disrupted in the disease. Critical cell-types for a biological function are defined as clusters of similar cells most “active” or “involved” in that biological function. In this paper, we provide a formal problem definition for the critical cell discovery problem using the recently introduced local correlation concept, and show that the proposed problem is intractable in theory. We propose a novel method, MoToCC (Module To Critical Cell-types), to find sets of similar cells with local correlated gene expression activity for input modules. We evaluated MoToCC on four neurodevelopmental disorder modules using single-cell expression data from the developing human cortex. Finally, we demonstrate that the objective value returned by MoToCC for the tested modules is an acceptable approximation to the optimal solution. Overall, our work provides a valuable tool for studying critical cell-types and their role in disease mechanisms, which could lead to the development of more effective targeted therapeutics. The MoToCC package is available at https://github.com/jchow32/MoToCC

## 1 Introduction

The advent of single-cell and single-nucleus RNA-sequencing (scRNA-seq and snRNA-seq) technologies has revolutionized the study of cellular decomposition in tissues by providing researchers with unbiased tools. These technologies enable the discovery of novel cellular subtypes and the genes that are differentially expressed in each of these subtypes. The identification of these subtypes has the potential to reveal the underlying etiology of complex diseases and common traits [1, 2]. Recent single-cell technologies have allowed the exploration of molecular mechanisms in a wide range of biological systems, including tumor cells and their associated microenvironments, as well as previously uncharacterized neuronal subtypes [3, 4]. Single-cell analysis enables the dissection of cellular heterogeneity and the identification of specific molecular targets for drug intervention, populations of cells with coordinated expression relevant to common biological functions, and the origins of disease pathogenesis, in contrast to bulk RNA-seq analysis [5, 6, 7]. Thus, single-cell and single-nucleus RNA sequencing have opened up new avenues for understanding the molecular basis of disease and for developing targeted therapies.

Genetic modules are groups of genes with similar biological functions that are distinct from other modules. These modules usually consist of co-expressed genes that are highly connected in protein-protein interaction networks. A genetic module that its disruption causes a specific disease is denoted as disease genetic module. Disease genetic modules often exhibit a significant enrichment of genetic variants or epigenetic modifications in affected cases compared to controls. Several methods for genetic module discovery have been developed, including those using gene expression, gene co-expression networks, protein-protein interaction networks, signaling pathways, and GO-terms as biological signals [8, 9, 10, 11]. The maturity of these methods and the ability to test modules in vitro and in vivo have led to the discovery of a growing list of disease genetic modules. To better understand the functional impact of a disease genetic module, it is crucial to identify the specific cell types in which the disease module is most active, and to understand how its disruption contributes to the disruption of critical cell types.

Previously proposed ideas for predicting the critical cell-types of certain diseases have involved linking the expression of modules to a specific defined tissue or cell-types [12, 13]. The simple approach of selecting the critical cell-type based on the highest expression of the input module assumes that the set of cell-types is static and predefined. In reality, the molecular profiles of individual cells are highly heterogeneous, thereby complicating the absolute assignment of a cell to any given cell-type or subtype.

In this paper, we propose a formal framework for the discovery of critical cell-types associated with a module. We have developed software named Module To Critical Cell-types (MoToCC), a novel linear programming approach that returns a subset of cells that selectively express module genes for given gene expression data. Users may vary the desired maximum number of cells to return as a solution, permitting the user to visualize relevant, distinct groups of cells that have similar expression levels at different scales of resolution. MoToCC source code and associated scripts are freely available at https://github.com/jchow32/MoToCC.

## 2 Problem Formulation

In this section, we will first provide an informal intuition about the critical cell discovery problem, next provide a formal problem definition, and finally give an NP-completeness proof of the proposed problems. We define a critical cell-type for a given genetic module as a set of cells that are the main target and impacted by the biological function of the input genetic module or pathway. Such approaches also provide a tool for finding the associations between human diseases, cell-types, and cellular processes by integrating various biological signals [14]. The level of expression of the genes in an input module can be considered as one factor in selecting the critical cell-types for an input module, however, we propose that the most important factor is the whole functional activity of the module that determines the critical cells. This is similar to utilizing gene co-expression (correlation between the expression of two genes) that has proven very effective in finding genetic modules.

Our formulation is motivated by the recently introduced concept of “local correlation” [12]. The local correlation between two genes is defined by considering their expression over all input cells, denoted by set *C*, available in the single-cell data [12]. Formally, they define local correlation statistics between two genes *p* and *q* as *h*_*p,q*_ = ∑_*i,j*∈*C*_ *w*_*i,j*_ *×* (*p*_*i*_*q*_*j*_ + *p*_*j*_*q*_*i*_), where *w*_*i,j*_ is the similarity score between cells *i* and *j*, and *p*_*i*_, *q*_*i*_, *p*_*j*_, *q*_*j*_ are the normalized and scaled expressions of the genes *p, q* in respective cells *i* and *j*. Note that the scaled and normalized gene expression guarantees that for every gene *p*, the *E*[*p*_*i*_] = 0 and *V ar*[*p*_*i*_] = 1 over all cells in *C*. Here, we propose to extend the definition of pairwise local correlation to a local correlation of a genetic module *M* and a set of cells *S ⊂ C* as

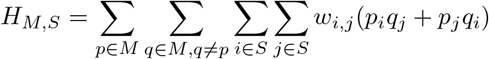

We are now ready to give formal problem definitions:

- First, we formally define the discovery of the most locally correlated cells (MLCC) problem. Assume a genetic module *M*, normalized single-cell gene expression, and a user-defined parameter *k* are provided. The objective of the MLCC problem is to select a subset cells *S ⊂ C* (where |*S*| ≤ *k*) to maximize *H*_*M,S*_.
- Next, we formally define the discovery of the critical cell-type problem as finding a subset cells *S ⊂ C* (where |*S*| ≤ *k*) to maximize *H*_*M,S*_ which are also “similar” based on available single-cell expression data. For the similarity constraint we use the condition that selected cells have to be strongly connected in the K-NN similarity graph. Note that the K-NN similarity graph is built using expression of all the genes [15].

**Theorem 1**. *The MLCC problem is NP-complete*.

*Proof*. We provide a reduction from the standard subset sum problem to the MLCC problem. Given a set of numbers, *X*, the decision version of the subset problem asks if there exists a nonempty subset of these numbers where their summation is equal to zero. We construct an instance of finding the most locally correlated cells problem using module *M* and set of cells *C*, i.e. finding the set *S ⊂ C* that maximizes the *H*_*M,S*_ given the module *M* and cells *C*, using numbers in set *X* as follows. We first assume module *M* consists of only two genes *p* and *q* (i.e., *M* = *{p, q}*), and for any number *x* ∈ *X* we consider a cell in set *C* with gene expression *x* and −*x* assigned to genes *p* and *q* respectively (i.e., *p*_*x*_ = *x, q*_*x*_ = −*x*). Note that total number of cells in this instance of the MLCC problem is equal to cardinality of *X* (i.e., |*C*| = |*X*|). We also assume the value *w*_*i,j*_ = 1 for every pair of cells *i* and *j*. Thus, solving this instance of the MLCC problem is finding a set of cells *S ⊂ C* that maximize *H*_*M,S*_ = ∑_*i*∈*S*_ ∑_*i*∈*S*_ (*p*_*i*_ *q*_*j*_ + *p*_*j*_ *q*_*i*_) = ∑_*i*∈*S*_ *p*_*i*_ ∑_*i*∈*S*_ *q*_*j*_ + ∑_*i*∈*S*_ *q*_*i*_ ∑_*i*∈*S*_ *p*_*j*_. Now, based on our instance construction assumption, we have ∑_*i*∈*S*_ *p*_*i*_ = ∑_*i*∈*S*_ *q*_*j*_. Thus *H*_*M,S*_ = − 2(∑_*i*∈*S*_ *p*_*i*_)^2^ ≤ 0. Finally, we conclude that a solution to this instance of the MLCC problem would also solve the decision version of the subset sum problem give set of numbers *X*.

### Cells with up-regulation activity of module *M*

Note that, in theory the above formulation the solution can produce a set of cells that the input module *M* is down-regulated in them as optimal solution. However, in practice we are mostly interested in critical cells that the input module *M* is up-regulated in them. Thus, we limit the above formulation by imposing an additional constraint such that module *M* is up-regulation in the selected cells.

We next propose a linear programming formulation for solving the MLCC problem, and then use this formulation with heuristic rounding to solve the critical cell-types problem.

## 3 Methods

We propose a two step framework, MoToCC, to discover the most critical cells given the module *M*. The inputs to MoToCC consist of a genetic module *M*, normalized single-cell gene expression, and a user-defined maximum number of critical cells *k*. The similarity matrices, Shared Nearest-Neighbor (SNN) and K nearest-neighbor (K-NN) graphs, are built using the normalized single-cell gene expression [16].

We first provide a linear programming (LP) formulation for solving the MLCC problem with a simple rounding solution of real values returned for the LP. Next, we will utilize the real value solution from the linear programming proposed for the MLCC problem to find the critical cell-types by imposing a cell-cell similarity constraint during the rounding step. The cell-cell similarity constraint in the critical-cell problem is that selected cells induced on the K-NN graph have to be a strongly connected component.

### 3.1 Linear programming formulation for MLCC problem

For solving the MLCC problem, we represent each cell as a node in a graph and for every pair of nodes *i* and *j*, we define a weight *w*_*i,j*_ indicating the cell-to-cell similarity as retrieved from the SNN graph. Given the genes in the input module *M*, we pre-calculate for every pair of cells *i* and *j* the score *z*_*i,j*_ =∑_*p*∈*M*_ ∑_*q*∈*M,p*≠*q*_ *w*_*i,j*_ (*p*_*i*_ *q*_*j*_ + *q*_*i*_ *p*_*j*_) where *p*_*i*_, *p*_*j*_, *q*_*i*_, and *q*_*j*_ represent the normalized expression of genes *p* and *q* (∈ *M*) in cells *i* and *j* respectively. The objective of the MLCC problem is to select a maximum of *k* cells such that the summation of *z*_*i,j*_ for every pair of cells selected is maximized. We note that the MLCC problem is NP-complete (Theorem 1) and thus we provide a heuristic solution for solving it.

We first introduce the linear programming (LP) relaxation of the MLCC formulation and use a simple rounding approach to get the final solution. For each cell/node *i* we define variable *y*_*i*_ to indicate the selection of that cell/node. For every pair of cells/nodes (*i, j*) we also define the variable *x*_*i,j*_ to indicate that both pairs of nodes *i* and *j* are selected (i.e., *y*_*i*_ = 1 and *y*_*j*_ = 1). We provide a linear programming formulation for the relaxed version of the MLCC problem as follows:

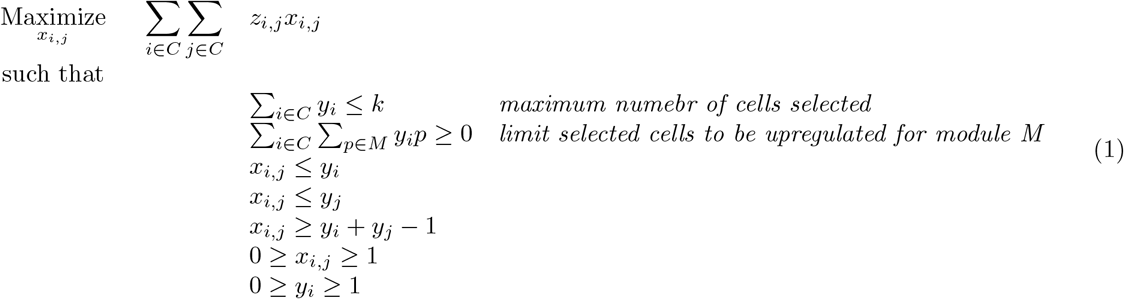

#### Rounding of MLCC solution

The proposed real value solution to the LP needs to be rounded to an integer solution indicating selected cells. As we are constrained by the number of cells we can select to be ≤ *k*, the rounding we propose is to select the top ≤ *k* cells with the highest calculated *y*_*i*_ score. Finally, the associated objective function values for cells in the LSCC are returned as the final solution.

### 3.2 Critical cell-types discovery problem

We propose a heuristic for predicting the critical cell-types that builds on the MLCC linear programming formulation. We utilize the same LP as the MLCC problem, however, during rounding we impose the constraint that selected cells need to be strongly connected in the K-NN graph. First, we select the initial solution returned by linear programming of cells with *y*_*i*_ *>* 0, which is referred to as potential “candidate cells”. Next, among the candidate cells, the largest strongly connected component (LSCC) in the K-NN graph is rounded to 1 and returned as the final solution. We utilize the largest strongly connected component (LSCC) function implemented in the networkx library (version 1.11) [17]. While, we don’t have a asymptotic approximation factor for this proposed rounding approach, we can experimentally calculate a lower bound on the approximation factor for various input modules using this solution (see section 4.4).

### 3.3 Implementation notes

For solving the linear programming we used the academic version of IBM ILOG CPLEX Optimizer [18]. To maximize the objective function (Section 2.2), edge weights (*z*_*i,j*_) are calculated between cells that have non-zero similarity (*w*_*i,j*_) as per the SNN graph. If pruning (–prune) is enabled, only edge weights outside of one standard deviation from the mean edge weight are retained. For a module, edge weights only need to be calculated once. Thus, if the user wishes to run MoToCC using varied k, the quickstart parameter (–quickstart) can be enabled to load edge weights previously calculated by MoToCC using the same module.

### 3.4 Return and refinement of initial solution

MoToCC returns the silhouette score associated with a 2D t-SNE dimensionality reduction and K-means clustering (K=2) of cells in the LSCC. Given multiple silhouette scores for varied k, breakpoints at which distinct groups of cell-types are selected can be viewed. To visualize the selected cells of the refined solution compared to unselected cells (via 2D t-SNE, 3D t-SNE, and UMAP) and to plot silhouette scores versus varied k, additional scripts and their usage are described at https://github.com/jchow32/MoToCC.

## 4 Results

### 4.1 Data

We evaluated the performance of the MoToCC pipeline on scRNA-seq gene expression datasets from the developing human cortex brain [13]. To analyze these datasets, we used four modules previously identified in studies of neurodevelopmental disorder: two NDD modules (M1 NDD and M2 NDD) that were discovered using *de novo* variants [19], an epilepsy module (M SCN1A) centered on the seed gene *SCN1A* [20], and an autism module (ASD mod13 Group1) enriched in chemical synaptic transmission genes [21]. We obtained and normalized the data using the standard Seurat pipeline.

### 4.2 Overview of experimental analysis

We compared the critical cells identified by MoToCC to those identified by Seurat and examined their agreement with known biological functions and cell subtypes associated with each module. To evaluate the statistical significance of critical cells selected per module, we compared the objective function value of the selected cells to that of randomly selected modules of the same size. To ensure that the gene expression distribution of the randomly selected modules was similar to that of the input module, we binned all genes into 10 bins based on their average expression across all cells and randomly selected the same number of genes from each bin as in the original input module. Finally, we compare the MoToCC results against a naïve heuristic approach that selects the subset of cell clusters found by Seurat [16] with the highest average gene expression, denoted as *h*_*S*_ - i.e., the average expression of module genes in selected cells S (see section 4.5).

The main results in support of the critical cells predicted by MoToCC:

- The critical cells predicted by MoToCC for all test modules are enriched in cell types associated with the disrupted module.
- For almost all the *k* values, MoToCC returns a significantly higher objective score using the real genetic module compared to randomly selected modules of the same size and similar average gene expression.
- MoToCC outperforms the naïve heuristic approach in selecting cell types with higher average gene expression for the input module.

### 4.3 Neurodevelopmental disorders critical cell-types

We investigated the critical cell-types impacted in NDDs using four different genetic modules previously reported [19, 20, 21]. For neurodevelopmental disorder modules, we consider two NDD modules found using MAGI [19], the epilepsy module found using MA<u>GI-S[20]</u> with *SCN1A* as a seed gene, and a recent autism module found [21] respectively denoted as M1 NDD, M2 NDD, M SCN1A and ASD mod13 Group1.

The M1 NDD module (80 genes) is significantly enriched in chromatin remodeling and the Wnt pathway, while the M2 NDD module (19 genes) is significantly enriched in chemical synaptic transmission and long-term potentiation. The module M SCN1A (36 genes) is significantly enriched in non-synonymous *de novo* mutations from epilepsy cohorts, known epilepsy genes, and pathways such as long-term potentiation, chemical synaptic transmission, and regulation of neurotransmitter activity. The ASD mod13 Group1 module (53 genes) was discovered via hierarchical clustering of a topological protein interaction network and was enriched in genes including *SYNGAP1, SHANK2*, and *SHANK3* and the synaptic transmission pathway.

The critical cells returned by MoToCC for all the NDD modules had significantly larger initial and final total objective function value at all *k* compared to modules of the same size consisting of random genes selected to have similar average gene expression. This indicates that the associated predicted critical cells with the input module have potential biological significance.

#### 4.3.1 Neurodevelopmental disorder module - M1 NDD

The M1 module exhibits functional enrichment in the regulation of transcription, chromatin remodeling, and WnT pathway, rather than neurotransmitter secretion. However, CSEA [22] failed to identify any significant enrichment of the M1 module in any specific cell type, emphasizing the need to consider single-cell data and specialized methods to identify critical cells.

Silhouette scores can indicate disparate clustering and shifts in percent composition. In the case of the M1 NDD module, red points highlight the largest increases in silhouette score (see Figure 1). Two-dimensional t-SNE plots were also displayed for several *k* cell values (50, 500, 1000, 1500, and 2000), with cells selected by LSCC appearing in red. The silhouette score analysis showed that the *k* cells between 750 and 1250 exhibited the sharpest increase, and the selected cells were mainly progenitor cells (PgS, PgG2M, and IP). This finding aligns with recent reports that emphasize the crucial role of the WnT pathway and primary autism genes in this module, including *CHD8*, in neural progenitor cells [23, 24].

**Figure 1.**
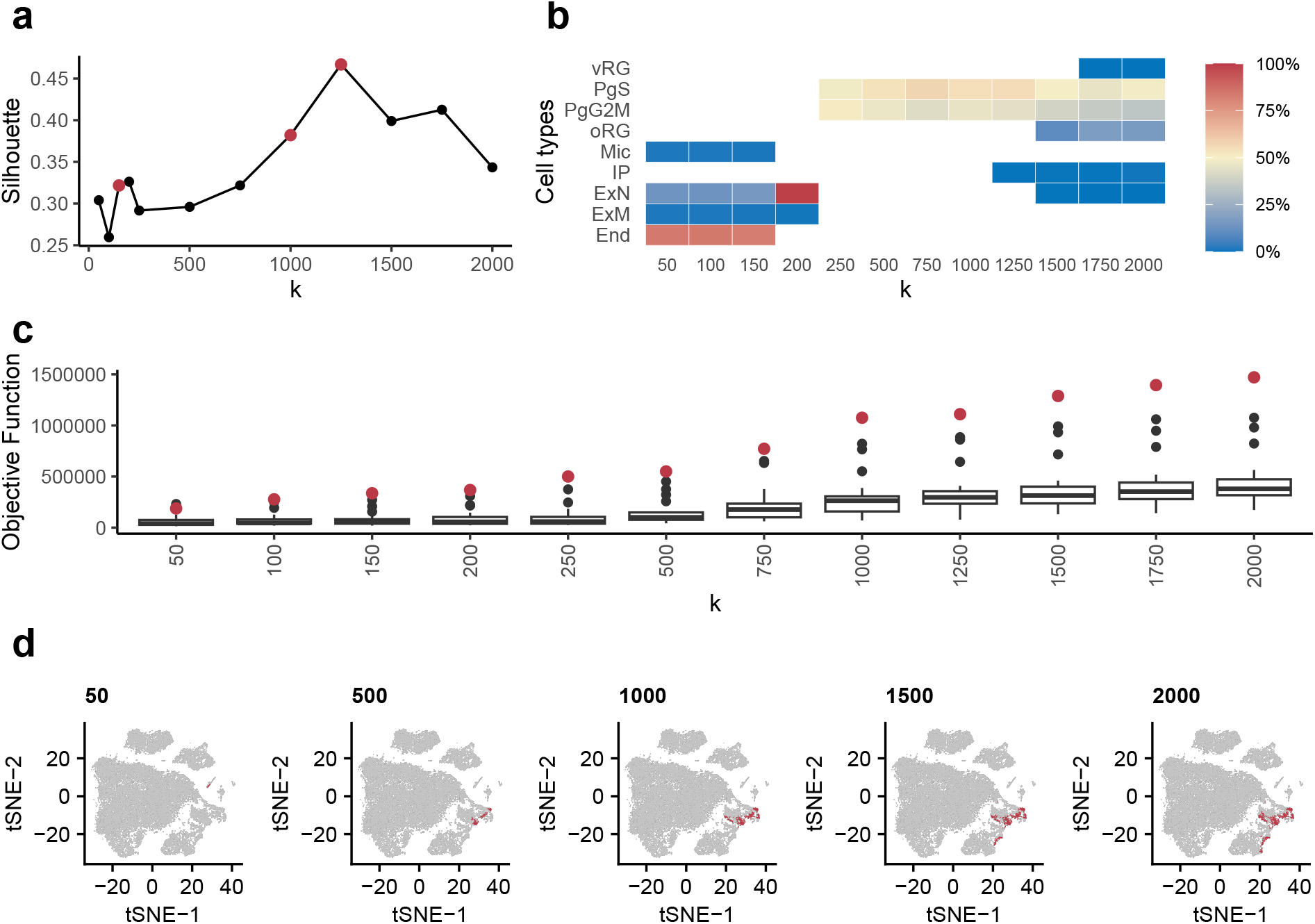
A) The largest increases in silhouette scores are highlighted in red. B) For the M1 module, selected cells from largest increase in silhouette score are primarily of the progenitor cell-type (PgS, PgG2M and IP). C) The final objective function of critical cells for the M1 module (red) are compared to values from 20 same-sized modules consisting of random genes (black). D) Two-dimensional t-SNE plots including selected critical cells (red) at *k* corresponding to the greatest increases in silhouette score (red title).

#### 4.3.2 Neurodevelopmental disorder module - M2 NDD

In this study, we also investigated M2 NDD, an NDD module identified using MAGI. The module M2 NDD is significantly enriched in synaptic transmission and long-term potentiation. Using MoToCC, we identified critical cells for M2 NDD across all *k* values, with the majority of the selected cells belonging to the excitatory deep layer 1 and 2 neurons (ExDp1, ExDp2) (Figure 2). Specifically, more than 67% of all cells of the ExDp2 cell-type were selected when k = 750, and the proportion of ExDp2 cells tends to increase as k increases for M2 NDD.

**Figure 2.**
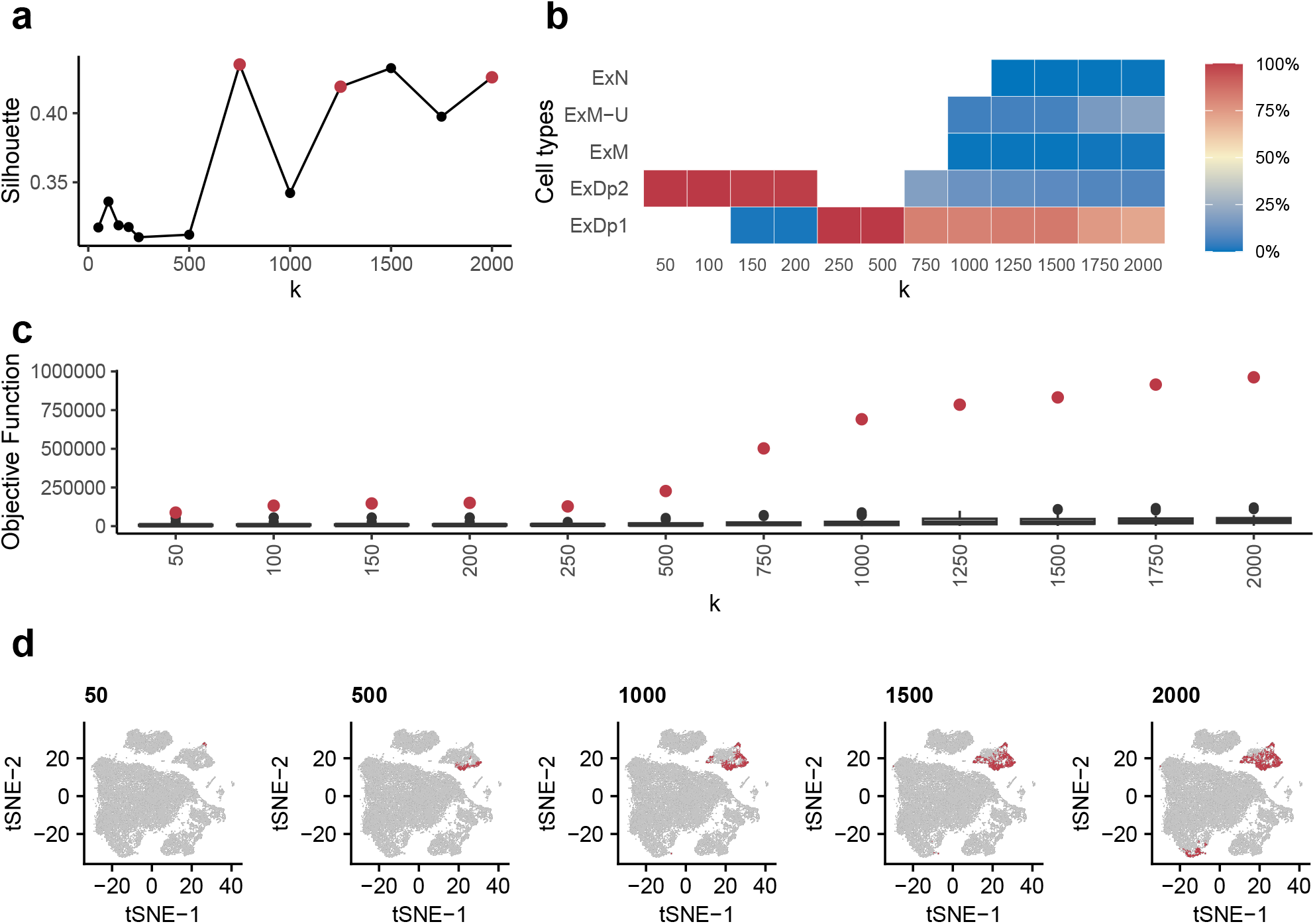
A) The largest increases in silhouette scores at k = (750; 1,250; 2,000) are highlighted in red. B) For the M2 NDD module, selected cells are primarily of the excitatory deep layer 1 and 2 (ExDp1, ExDp2) cell-types. C) The final objective function of critical cells for the M2 NDD module (red) are compared to values from 20 same-sized modules consisting of random genes (black). At all *k*, the corresponding final total objective function value of the M2 module is significantly greater than that of the randomized modules. D) Two-dimensional t-SNE plots including selected critical cells (red) at *k* corresponding to the greatest increases in silhouette score (red title). Cell-type abbreviations are provided in Supplementary table S1.

The elevated proportion of ExDp1 and ExDp2 cells in the M2 NDD module is consistent with our functional enrichment analyses, which reveal enrichment in synaptic transmission and regulation of neurotransmitter receptor and cation channel activity. Furthermore, this finding complements the increased expression levels of relevant genes such as *GABRB2, GRIN2B*, and *STXBP1* in ExDp2 cells. Consistent with these results, CSEA [22] also highlights the relevance of M2 genes in deep cortical neurons.

Taken together, our results suggest that the critical cells in M2 NDD are primarily excitatory deep layer 1 and 2 neurons, which are enriched in synaptic transmission and associated regulatory mechanisms. These findings provide valuable insights into the molecular and cellular mechanisms underlying neurodevelopmental disorders, and may help to inform the development of new therapeutic strategies.

#### 4.3.3 Epilepsy module - M SCN1A

The M SCN1A and M2 NDD modules display similar functional enrichment due to the large proportion (*>* 30%) of shared genes among modules. Like M2 NDD, critical cells of the M SCN1A module are primarily labeled as excitatory deep layer neurons. At smaller values of k (*k <* 250) the M SCN1A module’s critical cells initially consist of most existing ExDp2 cells (Figure 3). The percent composition of critical cells for higher k values is mostly ExDp1 and ExDp2. This is in agreement to what is reported as importance of excitatory neurons for SNC1A deficit samples [25, 26].

**Figure 3.**
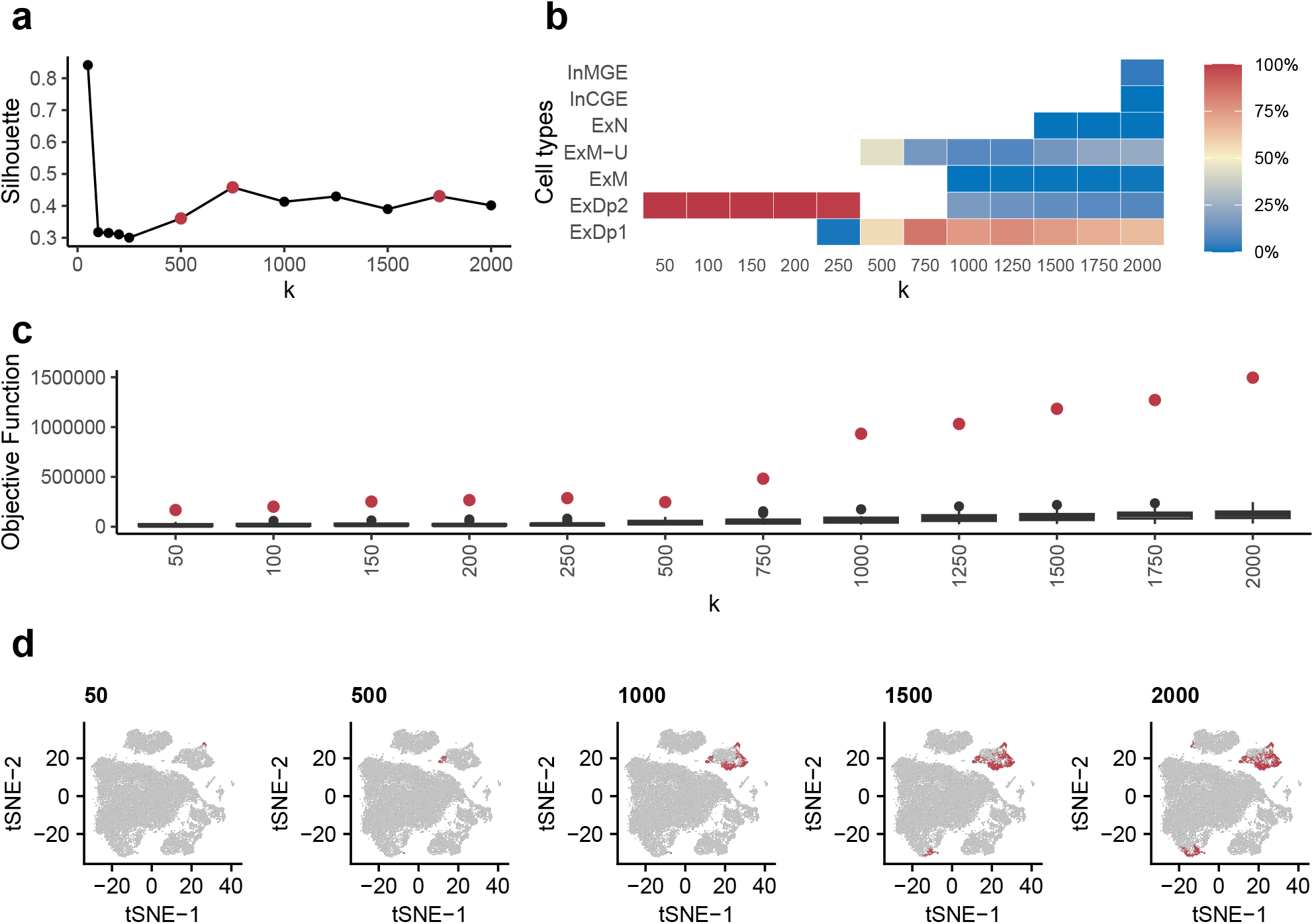
A) The largest increases in silhouette scores at k = (500; 7,50; 1,750) are highlighted in red. B) For the M SCN1A module, selected cells are primarily of the excitatory deep layer 1 and 2 (ExDp1, ExDp2) cell-types at lower k, followed by increased selection of the maturing excitatory upper enriched (ExM-U). C) The final objective function of critical cells for the M SCN1A module (red) are compared to values from 20 same-sized modules consisting of random genes (black). At all k, the corresponding final total objective function value of the M SCN1A module is significantly greater than that of the randomized modules. D) Two-dimensional t-SNE plots including selected critical cells (red) at k corresponding to the greatest increases in silhouette score (red title).

#### 4.3.4 Autism model (ASD Module 13, Group 1)

In a previous study, by Li et al. 2014 [21] found that the ASD mod13 Group1 module exhibited high expression specificity in the corpus callosum, a brain region mainly populated by oligodendrocyte progenitor cells (OPCs) and astrocytes. Unlike the M1 NDD, M2 NDD, and M SCN1A modules, ASD mod13 Group1 shares no common genes except for the seed gene *SCN1A* in M SCN1A. When using MoToCC to identify critical cell types at k=200 and 250, the algorithm selected only OPCs, which were followed by outer radial glial (oRG) and ventral radial glial (vRG) cells (see Figure 4). For k values below 150, MoToCC identified endothelial cells as the critical cell type, which showed the highest increase in silhouette score.

**Figure 4.**
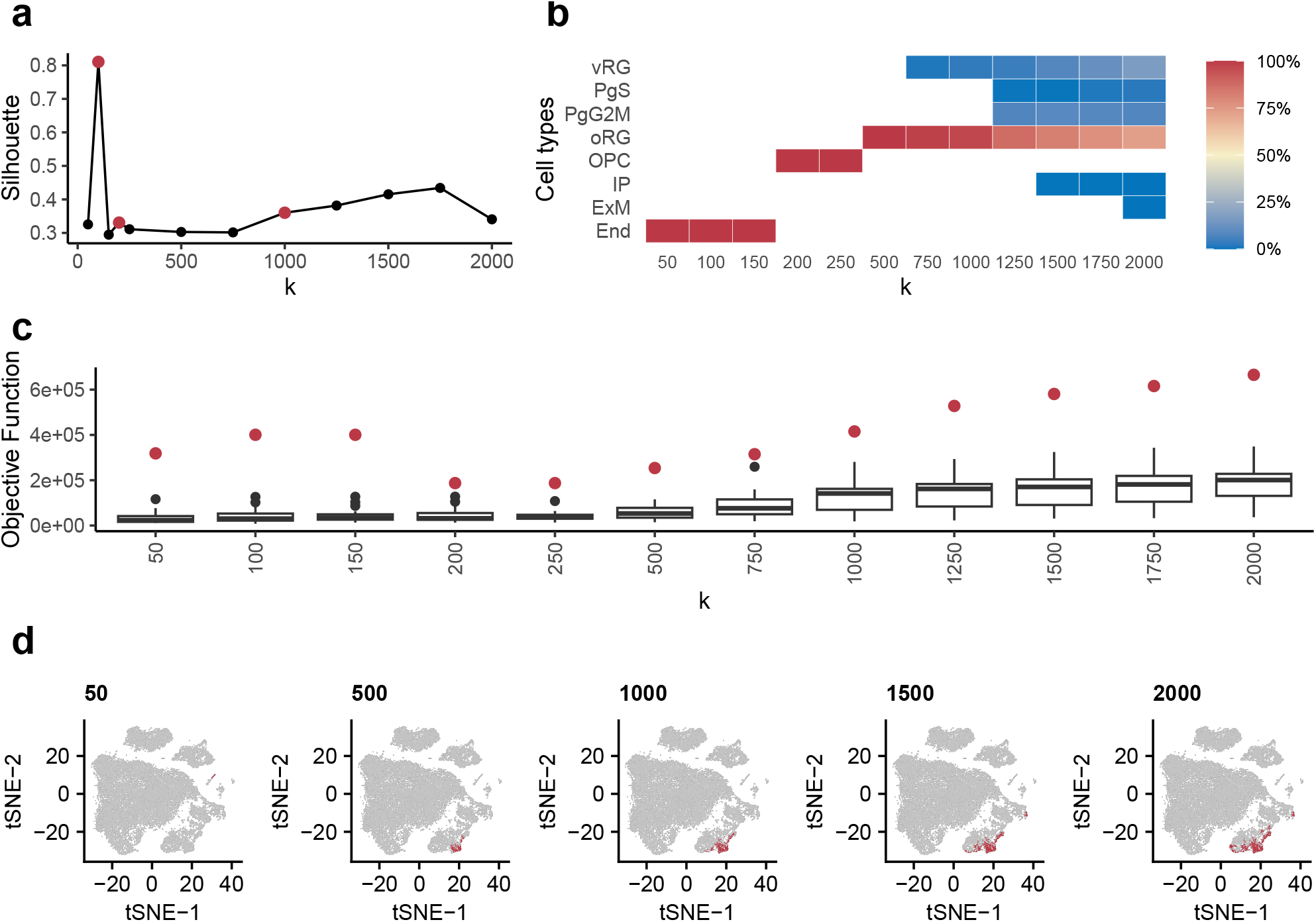
A) The largest increases in silhouette scores at *k* = (100; 250; 1,000) are highlighted in red. B) For the ASD mod13 Group1 module, selected cells for *k <* 150 are primarily endothelial cells and for *k >* 150 of the oligodendrocyte progenitor cell (OPC) cell-types. C) The final objective function of critical cells for the ASD mod13 Group1 module (red) are compared to values from 20 same-sized modules consisting of random genes (black). D) Two-dimensional t-SNE plots including selected critical cells (red) at *k* corresponding to the greatest increases in silhouette score (red title).

### 4.4 Estimation of approximation factor of critical cell discovery problem

We further investigate the empirical approximation factor of the proposed rounding heuristic for critical cell discovery using the four modules for various values of *k*. We calculated the ratio of final objective function value of *H*_*M,S*_ for selected cells over the objective function value returned by the linear programming as an empirical estimation of the approximation factor (Figure 5). Note that, as the objective value returned by the linear programming (using real value solution) is an upper bound on the true objective function we can conclude the ratio calculated an lower bound on the approximation factor (Figure 5). Based on observed ratio for the four different modules for various *k* it seems the approximation factor is constant and quite close to real solution (ratio is *>* 0.5 for most *k*).

**Figure 5.**
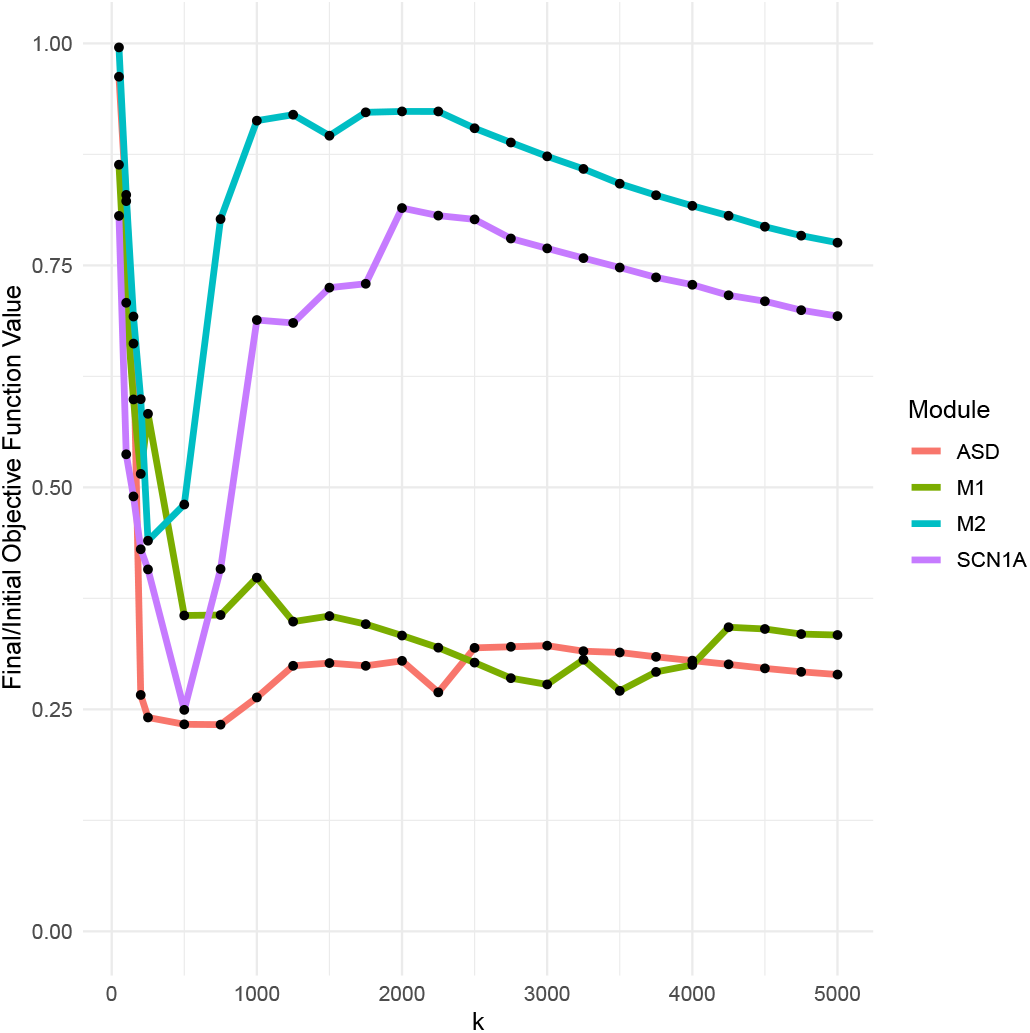
Ratio of final and initial objective function values for modules at varied k.

### 4.5 Comparison of MoToCC results against naïve approach

We provide a comparison of the MoToCC results against a naive approach, in which the cells labeled as belonging to the cell-type with the largest average normalized expression values (*h*_*s*_) were selected as a solution. While a ground truth in most cases is unknown, we compare the average expression (*h*_*s*_) for selected cells for the tested modules compared to naive selection of the cell-type associated with the largest *h*_*s*_ value (table 1). Note that, in all the five module while the MoToCC objective was not to maximize *h*_*s*_ value, yet it was able to achieve a higher *h*_*s*_ than the naive approach to select cells that maximize this score (table 1). For the M1

**Table 1:**
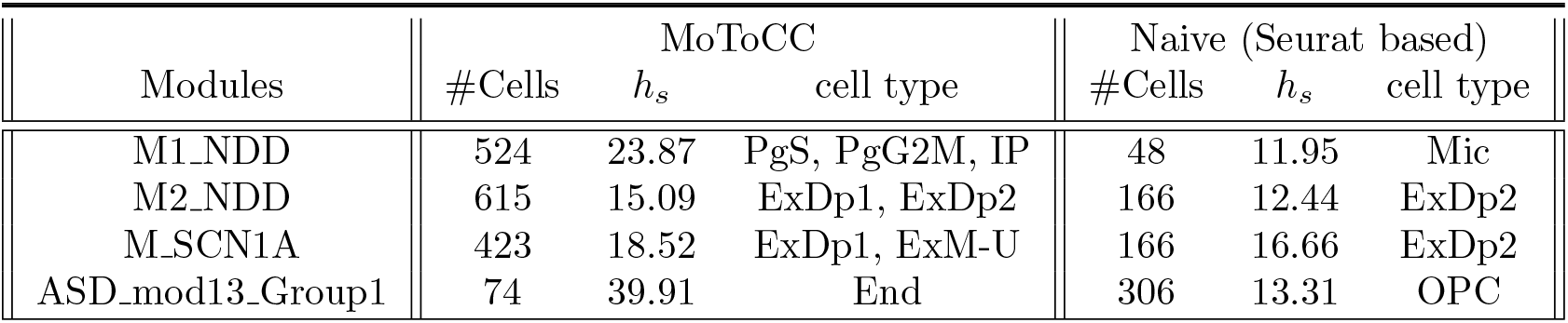
Comparison of average expression (*h*_*s*_) between MoToCC and naive (Seurat based) critical cell selection for selected *k* at elevated silhouette scores. Cell-type abbreviations are provided in Supplementary table S1.

## 5 Discussion

Understanding the critical cells affected by disrupted pathways in each disease is crucial for better understanding the etiology of diseases. To address this challenge, we have developed a novel approach called MoToCC that utilizes disrupted disease modules and single-cell expression data to identify the critical cells impacted in a disease associated with the input module.

MoToCC uses a linear programming approach to identify a set of cells that selectively express genes from a given genetic module. Specifically, MoToCC maximizes an objective function that considers both the correlated co-expression (local correlation) among module genes and cell-cell similarity. This optimization approach yields an initial solution, which is then refined using the derived K-nearest neighbor graph to select the largest strongly connected component (LSCC) among cells in the initial solution. Therefore, the selected critical cells in the LSCC consist of cells that share a high degree of local correlation in module genes and display similar overall gene expression profiles compared to all other cells in the dataset.

To assess the performance of MoToCC, we analyzed normalized single-cell gene expression data from the developing human cortex which were obtained from publicly available datasets [13]. We focused on four modules that are relevant to neurodevelopmental disorders (M1 NDD, M2 NDD, M SCN1A, and ASD mod13 Group1).

To identify the critical cells impacted by the disrupted pathways associated with each module, we used MoToCC to solve the linear programming problem and vary the upper bound *k* of the number of cells to return as a solution from 250 to 5,000 cells. By examining changes in silhouette score, we identified emphasized clusters of cells that were most relevant to the module and the targeted phenotype.

In general, at all *k*, the total objective function values associated with initial and final solutions tended to be significantly smaller for randomized modules of the same size as true modules, suggesting that cells were non-randomly selected according to their correlated gene expression.

The M1 NDD module is widely expressed in the human brain during neurodevelopment, with higher expression in the fetal brain. Functional analysis shows enrichment in chromatin remodeling, the Wnt pathway, and the Notch pathway. MoToCC identifies critical cells enriched in progenitor cells PgG2M and PgS, consistent with recent findings linking these cells to autism genes involved in chromatin remodeling and the Wnt pathway, such as *CHD8*.

M2 NDD and M SCN1A modules contain genes associated with synaptic transmission or neurotransmitter receptors, enriched in terms related to the regulation of neurotransmitter secretion. CSEA analysis shows selective expression in deep cortical neurons, crucial for functions such as memory formation and perception. For M2 NDD, ExDp1 and ExDp2 cell types were primarily selected, capturing over 67% of all ExDp2 cells. Example commands, guidelines, and associated scripts for pre-processing and data visualization are freely available at https://github.com/jchow32/MoToCC.

## 6 Acknowledgment

This project was partly funded by the BSF US-Israel Binational Science Foundation grant number 2021092 to FH and SS.

## 7 Supplementary Material

**Table S1:**
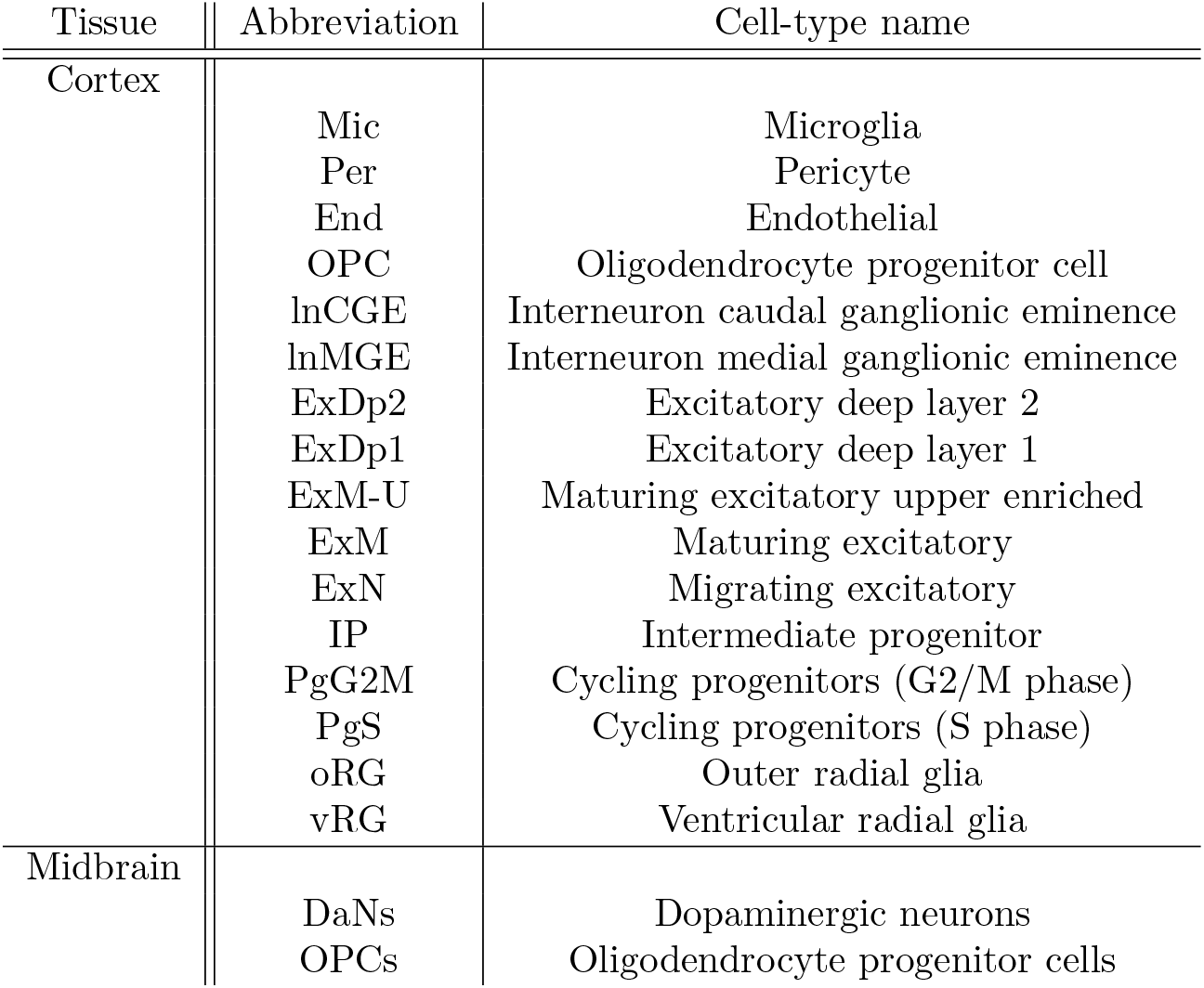
Cell-type abbreviations.

| Tissue | Abbreviation | Cell-type name |
| --- | --- | --- |
| Cortex | Mic | Microglia |
|  | Per | Pericyte |
|  | End | Endothelial |
|  | OPC | Oligodendrocyte progenitor cell |
|  | lnCGE | Interneuron caudal ganglionic eminence |
|  | lnMGE | Interneuron medial ganglionic eminence |
|  | ExDp2 | Excitatory deep layer 2 |
|  | ExDp1 | Excitatory deep layer 1 |
|  | ExM-U | Maturing excitatory upper enriched |
|  | ExM | Maturing excitatory |
|  | ExN | Migrating excitatory |
|  | IP | Intermediate progenitor |
|  | PgG2M | Cycling progenitors (G2/M phase) |
|  | PgS | Cycling progenitors (S phase) |
|  | oRG | Outer radial glia |
|  | vRG | Ventricular radial glia |
| Midbrain | DaNs | Dopaminergic neurons |
|  | OPCs | Oligodendrocyte progenitor cells |

